# Toward A Reproducible, Scalable Framework for Processing Large Neuroimaging Datasets

**DOI:** 10.1101/615161

**Authors:** Erik C. Johnson, Miller Wilt, Luis M. Rodriguez, Raphael Norman-Tenazas, Corban Rivera, Nathan Drenkow, Dean Kleissas, Theodore J. LaGrow, Hannah Cowley, Joseph Downs, Jordan Matelsky, Marisa Hughes, Elizabeth Reilly, Brock Wester, Eva Dyer, Konrad Kording, William Gray-Roncal

## Abstract

Emerging neuroimaging datasets (collected through modalities such as Electron Microscopy, Calcium Imaging, or X-ray Microtomography) describe the location and properties of neurons and their connections at unprecedented scale, promising new ways of understanding the brain. These modern imaging techniques used to interrogate the brain can quickly accumulate gigabytes to petabytes of structural brain imaging data. Unfortunately, many neuroscience laboratories lack the computational expertise or resources to work with datasets of this size: computer vision tools are often not portable or scalable, and there is considerable difficulty in reproducing results or extending methods. We developed an ecosystem of neuroimaging data analysis pipelines that utilize open source algorithms to create standardized modules and end-to-end optimized approaches. As exemplars we apply our tools to estimate synapse-level connectomes from electron microscopy data and cell distributions from X-ray microtomography data. To facilitate scientific discovery, we propose a generalized processing framework, that connects and extends existing open-source projects to provide large-scale data storage, reproducible algorithms, and workflow execution engines. Our accessible methods and pipelines demonstrate that approaches across multiple neuroimaging experiments can be standardized and applied to diverse datasets. The techniques developed are demonstrated on neuroimaging datasets, but may be applied to similar problems in other domains.

## 1 Introduction

Testing modern neuroscience hypotheses often requires robustly processing large datasets. Often the labs best suited for collecting such large, specialized datasets lack the capabilities to store and process the resulting images [1]. A diverse set of imaging modalities, including electron microscopy (EM) [1], array tomography [2], CLARITY [3], light microscopy [4], and X-ray microtomography (XRM) [5] will allow scientists unprecedented exploration of the structure of healthy and diseased brains. The resulting structural connectomes, cell type maps, and functional data have the potential to radically change our understanding of neurodegenerative disease.

Traditional techniques and pipelines developed and validated on smaller datasets may not easily transfer to datasets that are acquired by a different laboratory or that are too large to analyze on a single computer or with a single script. Prior machine vision pipelines for EM processing, for instance, have had considerable success [6, 7, 8, 9, 10]. However, these pipelines may require extensive configuration and are not scalable [8], may require proprietary software and have unknown hyperparameters [9], or are highly optimized for a single hardware platform [10].

In other domains, computer science solutions exist for improving algorithm portability and reproducibility, including containerization tools like Docker [11] and workflow specification such as the Common Workflow Language (CWL) [12]. Cloud computing frameworks enable the deployment of containerized tools [13], pipelines for scalable execution of python code [14], and reproducible execution [15]. Workflow management and execution systems such as Apache Airflow [16] and related projects such as Toil [17] and CWL-Airflow [18] allow execution of pipelines on scalable cloud resources. Despite the existence of these tools, a gap currently exists for extracting knowledge from neuroimaging datasets (due to the general lack of experience with these solutions as well as a lack of neuroimaging-specific features). We propose a solution that includes a library of reproducible tools and pipelines, integration with compute and storage solutions, and tools to automate and optimize deployment over large (spatial) datasets.

We introduce a library of neuroimaging pipelines and tools, Scalable Analytics for Brain Exploration Research (SABER), to address the needs of the neuroimaging community. SABER introduces canonical pipelines for EM and XRM, specified in CWL, with a library of dockerized tools. These tools are deployed using the workflow execution engine Apache Airflow [16] using Amazon Web Services (AWS) Batch to scale compute resources with imaging data stored in the volumetric database bossDB [19]. Metadata, parameters, and tabular results are logged using the neuroimaging database Datajoint [20]. Automated tools allow deployment of pipelines over blocks of spatial data, as well as end-to-end optimization of hyperparameters given labeled training data.

We demonstrate the use of SABER for three use cases critical to neuroimaging using EM, XRM, and light microscopy methods as exemplars. While light microscopy is commonly used to image cell bodies and functional activity with calcium markers, EM offers unique insight into nanoscale connectivity [21, 22, 23, 24], and XRM allows for rapid assessment of cells and blood vessels at scale [25, 5, 26]. These approaches provide complementary information and have been successfully used on the same biological sample [5], as XRM is non-destructive and compatible with EM sample preparations and light microscopy preparations. Being able to extract knowledge from large-scale volumes is a critical capability, and being able to reliably and automatically apply tools across these large datasets will enable the testing of exciting new hypotheses.

Our integrated framework is an advance toward easily and rapidly processing large-scale data, both locally and in the cloud. Processing these datasets is currently the major bottleneck in making new, large-scale maps of the brain — maps that promise insights into how our brains function and are impacted by disease.

## 2 Results

### 2.1 Pipelines and Tools for Neuroimaging Data

To address the needs of the neuroimaging community, we have developed a library of containerized tools and canonical workflows for reproducible, scalable discovery. Key features required for neuroimaging applications include:

- Canonical neuroimaging workflows specified in CWL [12] and containerized, open-source image processing tools
- Integration of workflows with infrastructure to deploy jobs and store imaging data at scale
- Tools to optimize workflow hyperparameters and automate deployment of imaging workflows over blocks of data

Building on existing tools, our framework provides an accessible approach for neuroimaging analysis, and can enable a set of use cases for the neuroscientist by improving reproducibility and reducing barriers to entry.

### 2.2 Standardized Workflows and Tools

While many algorithms and workflows exist to process neuroimagery datasets, these tools are frequently lab and task specific. As a result, teams often duplicate common infrastructure code (e.g., data download or contrast enhancement) and re-implement algorithms, when it would be faster and more reliable to instead reuse previously vetted tools. This hinders attempts to reproduce results and accurately benchmark new image processing algorithms.

In our framework, workflows are specified by CWL pipeline specifications. Individual tools are then specified by an additional CWL file, a container file, and corresponding source code. This ensures a modular design for pipelines and provides a library of tools for the neuroscientist.

Initially, we have implemented two canonical pipelines for EM and XRM processing. For EM, we estimate graphs of connectivity between neurons from stacks of raw images. Given XRM images, we estimate cell body position and blood vessel position. Each of these workflows is broken into a sequence of canonical steps. Such a step-wise workflow can be viewed as a directed, acyclic graph (DAG). Each step of a pipeline is implemented by a particular containerized software tool. The specific tools implemented in our reference canonical pipelines are discussed below.

#### 2.2.1 Cell Detection from X-ray Microtomography and Light Microcscopy

XRM provides a rapid approach for producing large-scale sub-micron images of intact brain volumes, and computational workflows have been developed to extract cell body densities and vasculature [5]. Individual XRM processing tools have been developed for tomographic reconstruction [27], pixel classification [28], segmentation of cells and blood vessels [5], estimation of cell size [5], and computation of the density of cells and blood vessels [5]. Running this workflow on a volume of X-ray images produces an estimate of the spatially-varying density of cells and vessels. Cubic millimeter-sized samples (100 GB) can be imaged, reconstructed, and analyzed in a few hours [5].

To implement a canonical XRM workflow, we define a set of steps: extracting subvolumes of data, classifying cell and vessel pixel probabilities, identifying cell objects and vasculature, merging the results, and estimating densities. Details on data storage and access can be found in the implementation section. We defined dockerized tools implementing a random forest classifier, a Gaussian Mixture Model, and a U-net [29] for pixel classification and the cell detection and vessel detection strategies [5]. These tools provide a standard reference for the XRM community, and modular replacements can be made as new tools are developed and benchmarked against this existing standard. Figure 1 shows this canonical workflow for XRM data, with each block representing a separate containerized tool. Also shown in Panel B is example output from running the pipeline, highlighting the resulting cell body positions and blood vessels.

**Figure 1:**
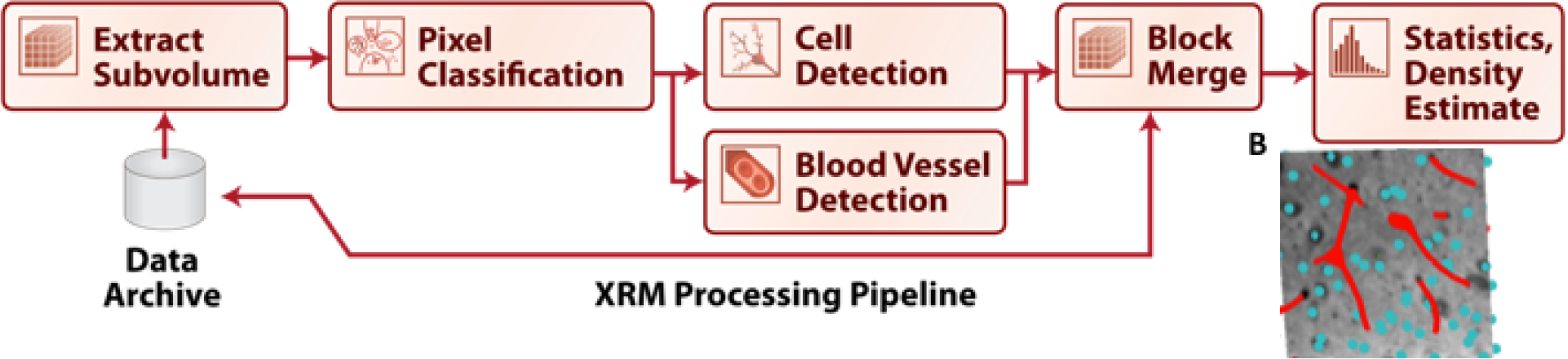
Workflow for processing XRM data to produce cell and vessel location estimates. Raw pixels are used to predict probabilities of boundaries, followed by detection of cell bodies and blood vessels. Finally, cell density estimates are created. Panel A shows the reconstruction pipeline, whereas Panel B shows a reconstruction of the detected cells and blood vessels in the test volume. Cells are shown as spheres and blood vessels as red lines.

These same tools can also be applied (with appropriate re-training) to detecting cell bodies from light microscopy data, such as from the Allen Institute Brain Atlas [4]. Here the same pipeline tools can be reused to detect cell bodies using the step for pixel classification followed by the step for cell detection. This result demonstrates the application of these tools across modalities and datasets to ease the path to discover.

#### 2.2.2 Deriving Synapse-level Connectomes from Electron Microscopy

Several workflows exist to produce graphs of brain connectivity from EM data [6, 10, 7], including an approach that optimizes each stage in the processing pipeline based on end-to-end performance [8]. However, these tools were not standardized into a reproducible processing environment, making reproduction of results and comparison of new algorithms challenging.

We have defined a series of standard steps required to produce brain graphs from EM images, seen in Figure 2. First, data is divided into subvolumes; cell membranes are estimated for each volume. Next, synapses are estimated and individual neurons are segmented from the data. After this, synaptic connections must be associated with neurons, and results merged together across blocks. Then a graph can be generated by iterating over each synapse to find the neurons representing each connection. Many tools have been developed for various sections of this pipeline, and a single tool may accomplish more than one step of the pipeline. Examples of tools for membrane segmentation include CNN [30] and U-nets [29] approaches. Synapse detection has been achieved using deep learning techniques and random forest classifiers [31, 32]. Neural segmentation has been previously done using agglomeration-based approaches [33] and automated selection of neural networks [9]. For our initial implementation of this workflow, we create CWL specifications and containerized versions of U-nets [29] for synapse and membrane detection, the GALA tool [34] for neuron segmentation, and algorithms for associating synapses to neurons and generating connectomes [8].

**Figure 2:**
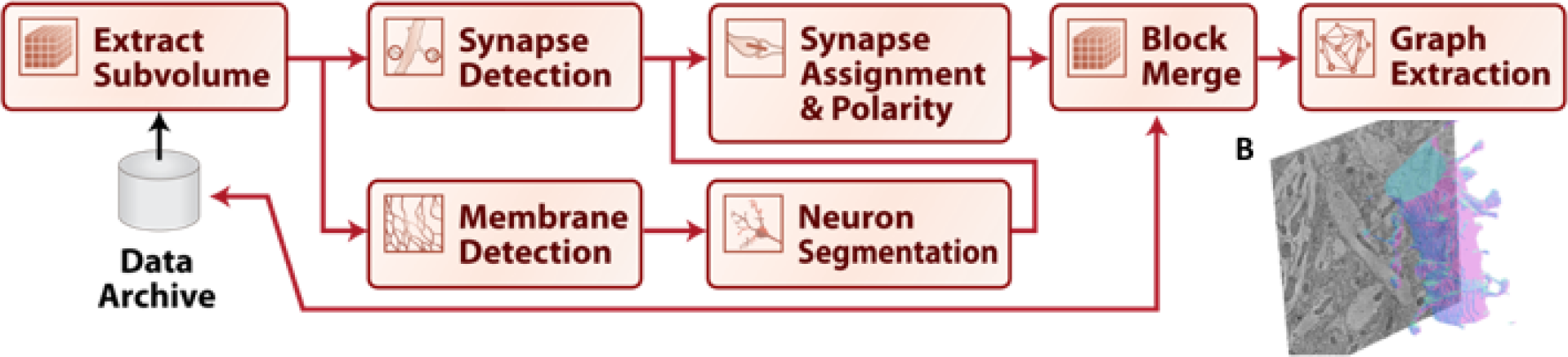
Canonical Workflow for Graph Estimation in EM data volumes. This workflow provides the ability to reconstruct a nanoscale map of brain circuitry at the single synapse level. The procedure of mapping raw image stacks to graphs representing synapse-level connectomes consists of synapse and membrane detection, segmentation of neurons, assignment of synapses, merging, and graph estimation. Panel A shows the reconstruction pipeline, and Panel B shows an example segmentation of a neuron from a block of data.

When creating this canonical pipeline for EM processing, our initial implementation goal is not to focus on pipeline performance in the context of reconstruction metrics. Rather, we aim to provide a reference pipeline for scientists and algorithms developers. For scientists, this provides an established and tested pipeline for initial discovery. For algorithm developers, this pipeline can be used to benchmark algorithms which encompass one or more steps in the pipeline.

### 2.3 Optimization and Deployment of Workflows

In addition to executing workflows the above workflows individually, it is critical for many use cases, such as in the neuroimaging applications detailed below, to launch workflows in parallel or sequentially, with modifications made to the tools and parameters. This controls functionality such as blocking data, running pipelines on each block, and merging results (i.e., a distribute-collect approach). It also enables end-to-end hyperparameter optimization and configuring tool selection. These are critical tools for discovery using neuroimaging pipelines, and we have implemented a python API to set workflow parameters, schedule jobs, and poll results. Deployment scripts enable rapid configuration and processing paradigms.

In order to perform hyperparameter optimization (Figure 3), our tools currently require a small volume of labeled training data (although are also exploring unsupervised methods). Because hand labeling is expensive, the team will host available ground truth data (e.g., [22] for EM), to facilitate scoring, using metrics such as precision-recall metric or f1-score. We pursue an optimization strategy that assumes a black-box workflow, avoiding assumptions such as differentiability of the objective function. These techniques are suitable for searching hyperparameters of various tools. To begin, we implemented a simple grid search, random search, and the adaptive search method shown in Figure 3, based on random resampling [35]. We plan to techniques such as sequential Bayesian optimization [36] and convex bounding approaches [37] to develop a library of readily available, proven techniques.

**Figure 3:**
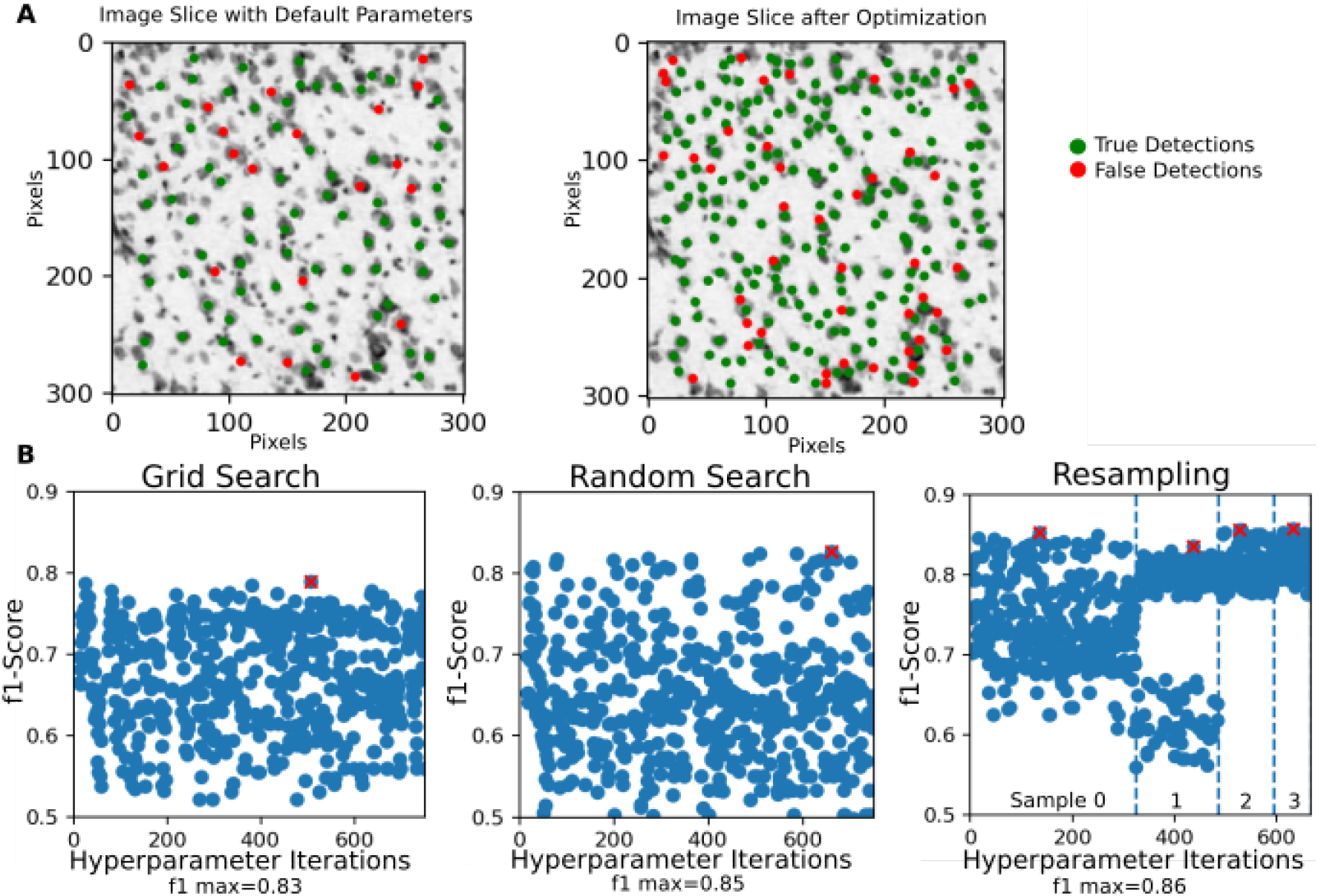
Use case of optimizing a pipeline for light microscopy data, comparing grid search, random search, and the random resampling approach described in the text. We demonstrate these tools on a light microscopy dataset, leveraging methods originally developed for XRM – showcasing the potential for applying tools across diverse datasets. The framework allows a user to easily compare the trade-offs of different approaches for a particular dataset. The maximum f1 score for each approach is marked with a red ’x’. Automating this process using SABER allows for rapid deployment and optimization.

## 3 Neuroimaging Use Cases

### 3.1 Use Case 1: Pipeline Optimization

When collecting a new neuroimaging dataset, it is often necessary to fine-tune or retrain existing pipelines. This is typically done by labeling a small amount of training data, which can often be labor intensive, followed by optimizing the automated image processing pipeline for the new dataset. These pipelines consist of heterogeneous tools with many hyperparameters and are not necessarily end-to-end differentiable.

Users can execute the optimization routines using a simple configuration file to specify algorithms, parameter ranges, and metrics. Figure 3 demonstrates the application of three algorithms for pipeline optimization. We choose the Allen Institute for Brain Science (AIBS) Reference Atlas [4] as a demonstration of generalization beyond EM and XRM datasets. In order to optimize the pipeline, this example optimizes over: the initial threshold applied to the probability map, size of circular template, size of circular window used when removing a cell from the probability map, and the stopping criterion for maximum correlation within the image. The user specifies the range of each parameter.

Our framework supports implementations of different optimization routines, such as random selection of parameters with resampling, as seen in Figure 3. Random selection of parameters often produces comparable results to grid search, and users may need to explore algorithms to find an approach that works well for the structure of their pipeline [35]. For the resampling approach, we initially choose parameters at random, and then refine search parameters by choosing new parameters near the best initial set, with the user setting a maximum number of iterations. Figure 3B shows a parameter reduction of twenty percent at each resampling, leading to a more efficient parameter search and improved performance. Using SABER, it is possible for a user to explore the trade-offs for a range of hyperparameter optimization routines.

### 3.2 Use Case 2: Scalable Pipeline Deployment

The second critical use case of interest to neuroscientists is the deployment of pipelines to large datasets of varying sizes. Datasets may be on the order of gigabytes or terabytes, as in XRM, to multiple petabytes, as in large EM volumes used for connectome estimation. SABER provides a framework for blocking large datasets, executing optimized pipelines on each block, then merging the results through a functional API. Given a dataset in a volumetric database, such as bossDB, our python scripts control blocking, execution, and merging. Results are placed back into a database for further analysis, or stored locally. An example of this use case for XRM data can be seen in Figure 4, and another example of this use case for extracting synapse-level connectomes can be seen in Figure 5.

**Figure 4:**
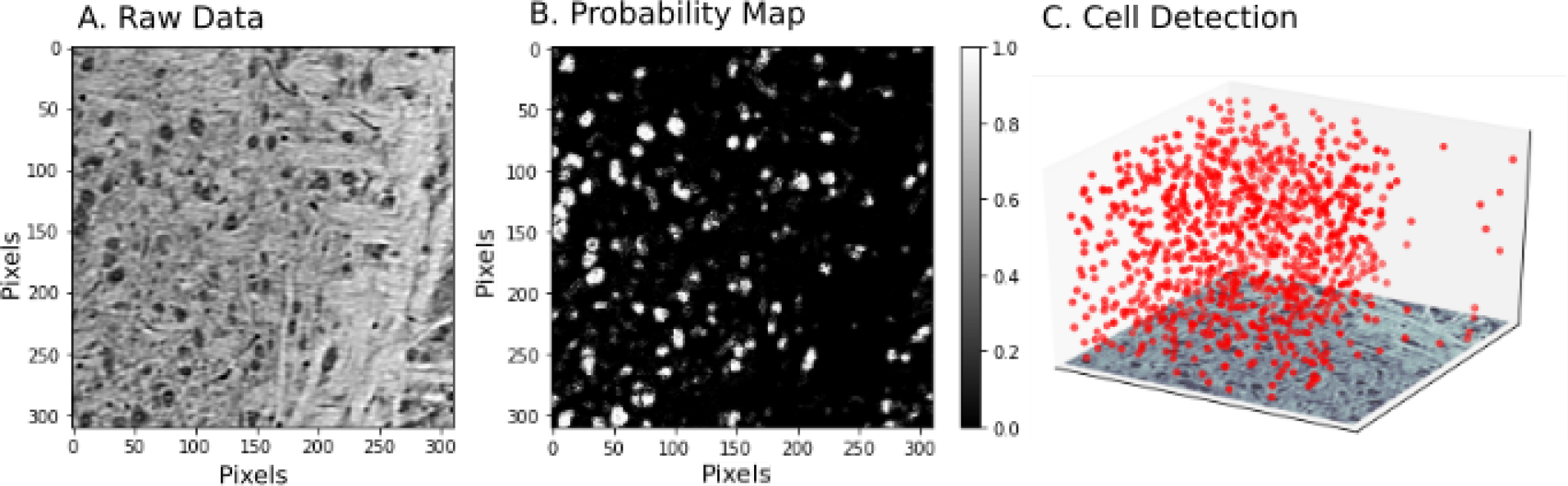
Example deployment of pipeline over spatial dataset, in this case cell detection in XRM data. An example slice of raw data can be seen in Panel A. The pipeline in Fig.1 was used to classify pixels (Panel B) and detect cells. From the cells, a three dimensional scatter plot of the positions of the cell centers was generated (Panel C).

**Figure 5:**
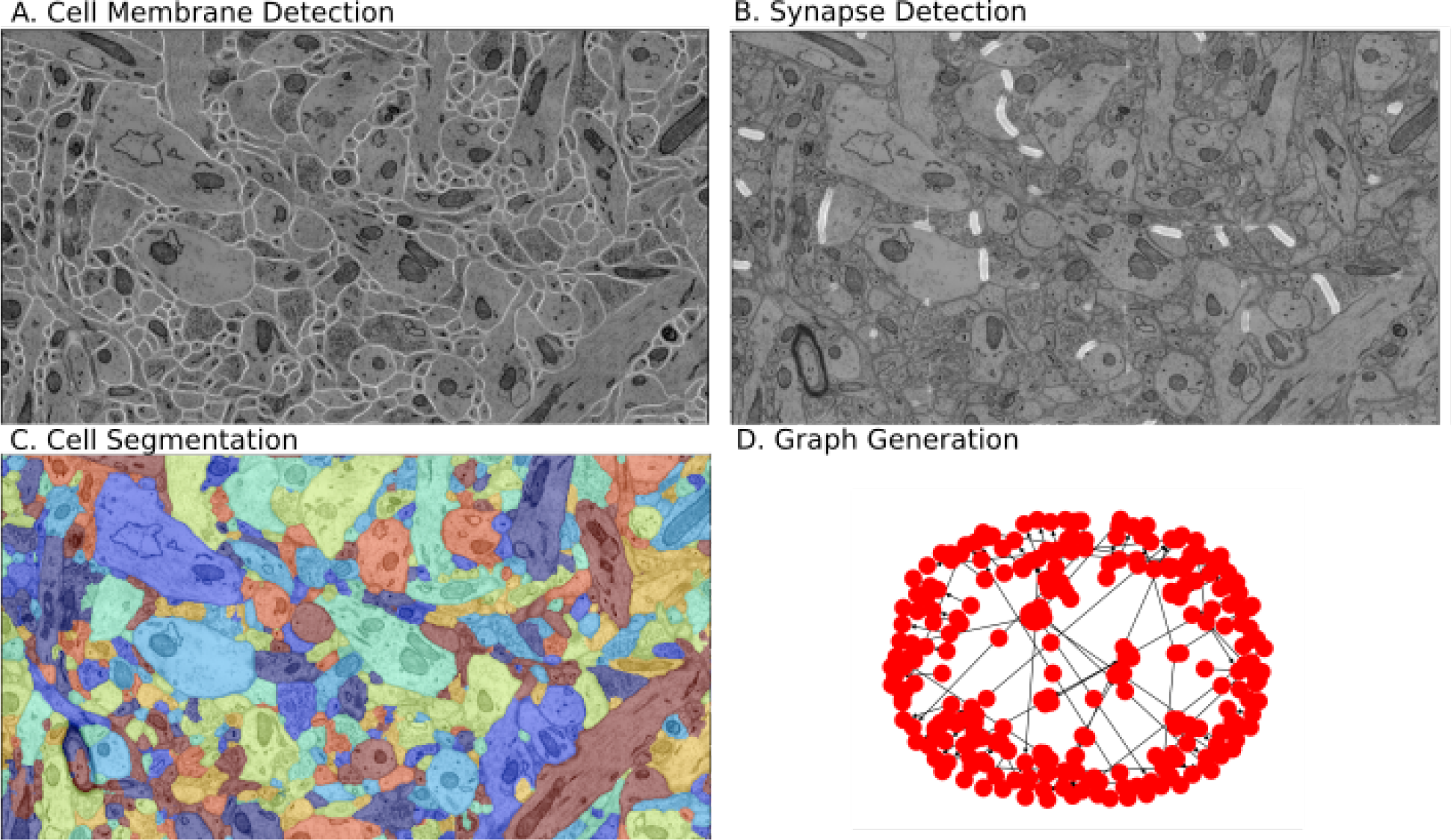
Example deployment of EM segmentation pipeline to extract graphical models of connectivity from raw images. The processing pipeline (Fig.1) consists of neural network tools to perform A) membrane detection and B) synapse detection. This is followed by a segmentation tool (Panel C). Finally, segmentation and synapses are associated to create a graphical model.

### 3.3 Use Case 3: Benchmarking Neuroimaging Algorithms

The third major use case applies to developers implementing new algorithms for neuroimaging datasets. Due to tools being written in a variety of languages for a variety of platforms, it has been difficult for the community to standardize comparison between algorithms. Moreover, it is important to assess end-to-end performance of new tools in a pipeline which has been properly optimized. Without this comparison, it is difficult to directly compare algorithms or their impact. Using the specified pipelines, a new tool may subsume one or more of these steps, with the specification defining the inputs and outputs. A new CWL pipeline can be quickly specified with the new tool replacing the appropriate step or steps. Hyperparameter optimization can be run on each example to compare tools, leveraging reference images and annotations for the pipelines provided in SABER.

## 4 Discussion

We have developed a framework for neural data analysis along with corresponding infrastructure tools to allow scalable computing and storage. We facilitate the sharing of workflows by compactly and completely describing the associated set of tools and linkages. Future enhancements will introduce versioning to track changes in workflows and tools.

Our goal is to establish accessible reference workflows and tools which can be used for benchmarking new algorithms and assessing performance on new datasets. Moving forward, we will encourage algorithm developers to containerize their solutions for pipeline deployment and to incorporate state-of-the-art methods. Through community engagement, we hope to grow the library of available algorithms and demonstrate large-scale pipelines which have been vetted on different datasets. We also hope to recruit researchers from different domains to explore how these tools apply outside of the neuroimaging community.

Prior solutions have taken different approaches to processing neuroimaging data. For example, the workflow execution engine LONI has been used for processing EM data [8], but requires extensive configuration and is not scalable to very large volumes. The SegEM framework [9] offers extensive features for optimizing and deploying EM pipelines, but is specifically focused on neuron segmentation from EM data and is tied to a MATLAB cluster implementation. Highly optimized pipelines can be deployed on a single workstation [10], which is ideal for proven pipelines as part of ongoing data collection, but is limited in developing and benchmarking new pipelines.

A limitation of our existing tooling is interactive visualization. Although we provide basic capabilities, additional work is needed to interrogate raw and derived data products and identify failure modes. We are extending open source packages, substrate [38] and neuroglancer [39], to easily visualize data inputs and outputs of our workflows and tools.

Scalable solutions for container such as Kubernetes [13] and general workflow execution systems like Apache Airflow [16] have provided the ability to orchestrate execution of containers at scale. These solutions, however, lack workflow definitions, imaging databases, and deployment tools to enable neuroimaging usecases. SABER builds on top of these technologies to enable neuroimaging use cases while avoiding the specialized, one-off approaches often used in conventional neuroimaging pipelines.

## 5 Potential implications

While our initial workflows focus on XRM and EM datasets, many of these methods can be easily deployed to other modalities like light microscopy [40], and the overall framework is appropriate for problems in many domains. These include other scientific data analysis tasks as varied as machine learning for processing noninvasive medical imaging data or statistical analysis of population data.

Code, demos, and results of the SABER platform are available on GitHub under an open source license, along with documentation and tutorials (see below). We make SABER available to the public with the expectation it will help to enable and democratize scientific discovery of large, high-value datasets, and that these results will offer insight into neurally-inspired computation, the processes underlying disease, and paths to effective treatment.

## 6 Methods

### 6.1 Existing Software Solutions

For small-scale problems, individual software tools and pipelines which are fully portable and reproducible have been produced (e.g., [41]), but this challenge has not yet been solved at the scale of modern EM and XRM volumes.

Many tools have become available for scalable computation and storage, such as Kubernetes [13] and Hadoop [42], which enable the infrastructure needed for running containerized code at scale. However, such projects are domain-agnostic and do not necessarily provide the features or customization needed by a neuroscientist. As scalable computation ecosystems, these solutions can be integrated as the backend for workflow management systems such as SABER.

Traditional workflow environments (e.g., LONI Pipeline [43], Nipype [44], Galaxy [45], and Knime [46]) provide a tool repository and workflow manager, but require connection to a shared compute cluster to scale. All of these systems rely on software that are installed locally on the cluster or local workstation, and can result in challenging or conflicting configurations that slow adoption and hurt reproducibility.

New frameworks for workflow execution have been developed, but solve only a subset of the challenges for neuroimaging. Boutiques [47] manages and executes single, command-line executable neuroscience tools in containers. Pipelines must be encapsulated in a single tool, meaning that coding is required to swap pipeline components. Dray [48] executes container-based pipelines as defined in a workflow script. While Dray contains some of the core functionality to execute container-based pipelines, non-programmers cannot easily use the system and it is limited in the types of workflows that are supported.

Similarly, Pachyderm [49], offers execution of containerized workflows but lacks support for storage solutions appropriate for neuroimaging as well as optimization tools needed for these neuroimaging pipelines. Workflow execution engines such as Toil [17] and CWL-Airflow [18] are closely related to SABER, providing light-weight python solutions for workflow scheduling. However, like Pachyderm, they lack the automation tools and storage scripts required by neuroimaging applications. The most closely related tool is Air-tasks [50], which provides tools to automate deployment of neuroimaging pipelines. Air-tasks, however, provides fewer capabilities to the user and does not support a common workflow specification or explicitly support optimization or benchmarking.

### 6.2 SABER

To overcome limitations in existing solutions, SABER provides canonical neuroimaging workflows specified in a standard workflow language (CWL), integration with a workflow execution engine (Airflow), imaging database (bossDB), and parameter database (Datajoint) to deploy workflows at scale, and tools to automate deployment and optimization of neuroimaging pipelines. Our automation tools include end-to-end hyperparameter optimization methods and deployment by dividing data into blocks, executing pipelines, and merging results (block-merge).

The core framework (called CONDUIT) is provided in a docker container to reduce installation constraints and increase portability (Figure 6). The core framework interfaces with scalable cloud compute and storage resources. A comparison of SABER/CONDUIT to existing solutions is seen in Table 1.

**Table 1:** Comparison of existing projects related to workflow execution of neuroimaging pipelines with green cells highlighting desirable features, yellow cells highlight partial implementations of desirable features, and red cells highlighting limitations. SABER delivers integrated containerized tools, a standardized workflow and tool description, and a volumetric database. It also provides tools for automating deployment over datasets by dividing into blocks (block-merge) and optimization of workflows. The most comparable tools are other workflow management systems such as CWL-Airflow, TOIL, Galaxy, and Air-tasks. Air-tasks provides similar capabilities, but lacks support for common workflow descriptions and tool optimization, and less flexibility for users. Similar projects such as TOIL, Galaxy, and CWL-Airflow lack neuroimaging specific features to enable the use cases described in Section 3. Scalable cluster systems, such as Kubernetes, provide essential functionality to deploy containers at scale, but need capabilities built to manage workflows and data movement and are complementary to workflow management systems such as SABER.

|  | **SABER/**<br>**CONDUIT** | **CWL-**<br>**Airflow** | **TOIL** | **Galaxy** | **Air-tasks** | **Kubernetes** |
| --- | --- | --- | --- | --- | --- | --- |
| Purpose | Workflow management system | Workflow management system | Workflow management system | Workflow management system | Workflow management system | Distributed Container Orchestration |
| Container Support | Yes | Yes | Yes | Yes | Yes | Yes |
| Workflow Description | CWL | CWL | CWL or WDL | Custom (CWL beta) | Custom python | N/A |
| Computational background | Novice-Expert | Novice-Expert | Novice-Expert | Novice-Expert | Intermediate-Expert | Expert |
| Installation | docker-compose | pip install | pip install | Install scripts | docker-compose | Cluster configuration |
| Cloud Support | AWS | Planned | Multiple cloud providers | Multiple cloud providers | Docker Infrakit | Multiple cloud providers |
| Volumetric Database | bossDB, DVID | None | None | None | Cloud Volume | N/A |
| Parallel Processing Model | Block-merge | None | None | None | Block-merge | N/A |
| Workflow deployment for neuroimaging | Yes | No | No | No | Yes | N/A |
| Workflow optimization for neuroimaging | Yes | No | No | No | No | N/A |
| Tool Benchmarking and Datasets | Yes | No | No | No | No | N/A |
| EM tool library | Yes | No | No | No | Yes | N/A |
| Tools for Other Modalities | Yes | No | No | No | No | N/A |

**Figure 6:**
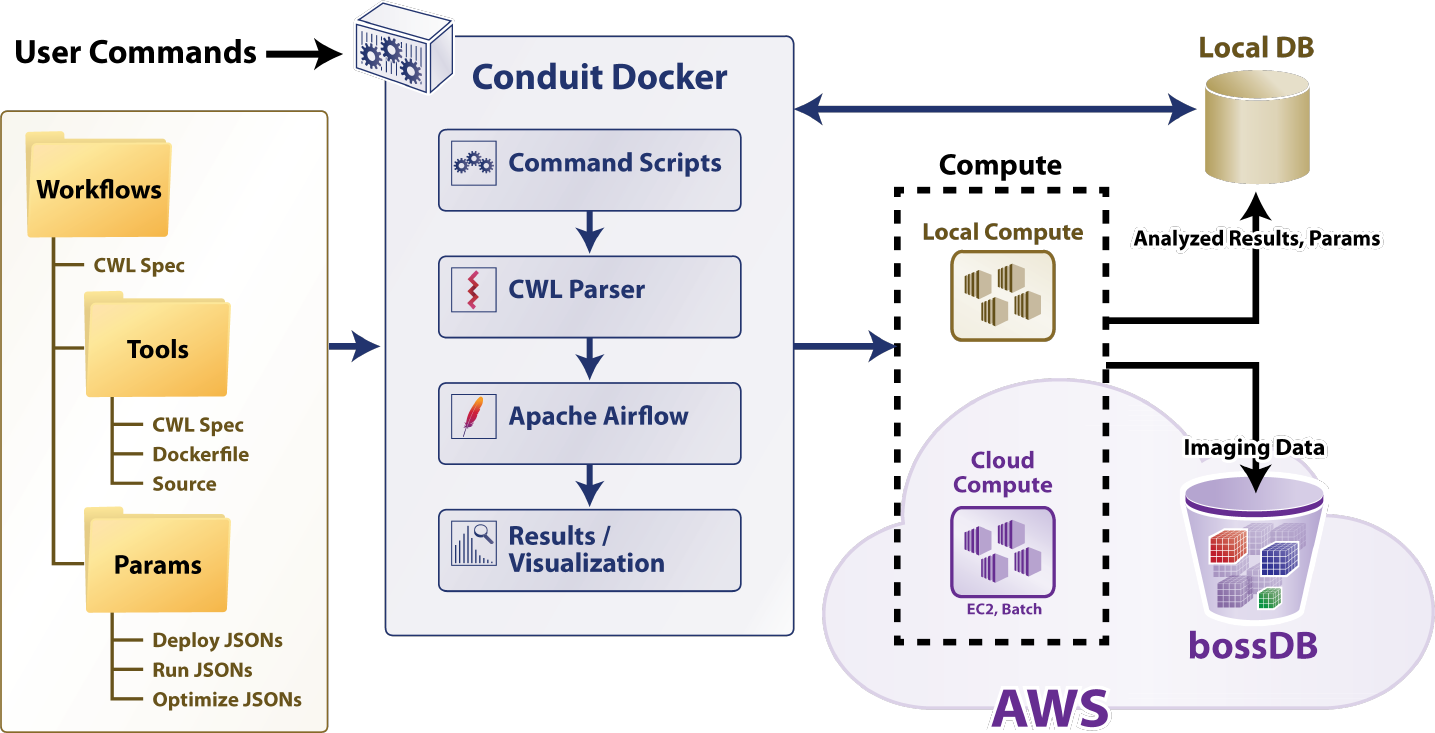
The architecture and components of SABER. Tools, workflows, and parameters for individual use cases (optimization, deployment) are captured in a file structure using standardized CWL specifications and configuration files. The core of the framework (called CONDUIT) is run locally in a docker container. CONDUIT consists of scripts to orchestrate deployment and optimization, a custom CWL parser, Apache Airflow for workflow execution, and tools to collect and visualize results. Containerized tools are executed locally or using AWS Batch for a scalable solution. The bossDB provides a solution for scalable storage of imaging data, and a local database is used for storing parameters and derived information.

In our CONDUIT framework (Figure 6 highlights the architecture of the system), workflows and tools are defined with CWL v1.0 specifications. Tools additionally include Dockerfiles and source code. Parameter files contain user-specified parameters for optimization and deployment of pipelines.

#### 6.2.1 Workflow Execution

The CONDUIT container shown in Figure 6 provides SABER with a managed pipeline execution environment that can run locally or scale using the AWS Batch service. Our custom command scripts and CWL parser generate DAG specifications for execution by Apache Airflow. We select Apache Airflow to interface with a cloud-based computing solution. As an example, we utilize the AWS Batch service, although Airflow can interface with scalable cluster solutions such as Kubernetes or Hadoop. The framework facilitates the execution of a batch processing (versus streaming) workflow composed of software tools packaged inside multiple software containers. This reduces the need to install and configure many, possibly conflicting software libraries.

##### 6.2.2 Cloud Computation and Storage

Large neuroimaging datasets are distinct from many canonical big data solutions because researchers typically analyze a few (often one) very large datasets instead of many individual images. Custom storage solutions [19, 51] exist, but often require tools, knowledge, and access patterns that are disparate from those used by many neuroscience laboratories. SABER provides tools to connect to specialized neuroimaging databases which integrate into CWL tool pipelines. We use intern [52] to provide access to bossDB and DVID and abstract data storage, RESTful calls, and access details. Workflow parameters, objective functions, and summary results such as graphs and cell densities can be stored using a DataJoint database [20] using a custom set of table schemas.

Some datasets, however, can be stored locally but are too large to process in memory on a single workstation. In addition to volumetric data stored in bossDB, SABER also supports local imaging file formats such as HDF5, PNG, or TIFF. As users share pipelines, they might wish to use a pipeline originally developed for data stored in one archive with that stored in another.

Modern cloud computing tools, such as AWS Batch or Kubernetes, allow large scale deployment of containerized tools on demand. The CONDUIT container schedules workflows using Apache Airflow, which supports execution using these scalable containerized tools. Workflows can also be executed using local resources for smaller jobs.

## 7 Availability of source code and data

The SABER framework is open source and available online:

- Project name: SABER
- Project home page: e.g. https://github.com/aplbrain/saber
- Operating system(s): Platform independent
- Programming language: Python, other
- Other requirements: Docker, AWS account (if scalable cloud computing required)
- License: Apache License 2.0

The source code for this project is available on GitHub, including code for tools and demonstration workflows. An extensive wiki documenting the repository is also hosted on github. The data are stored in a bossDB instance at https://api.bossdb.org.

## 7.1 Acknowledgements

We would like to thank the Apache Airflow and Common Workflow language teams for their open-source tools supporting reproducible workflows, as well as the research groups who produced our reference EM and XRM volumes for analysis. Research reported in this publication was supported by the National Institute of Mental Health of the National Institutes of Health under Award Number R24MH114799. The content is solely the responsibility of the authors and does not necessarily represent the official views of the National Institutes of Health.

